# Cellular and gene expression patterns associated with root bifurcation in Selaginella

**DOI:** 10.1101/2022.01.03.474808

**Authors:** Tao Fang, Hans Motte, Boris Parizot, Wouter Smet, Xilan Yang, Liam Walker, Maria Njo, George W. Bassel, Tom Beeckman

## Abstract

The roots of lycophytes branch through dichotomy or bifurcation, which means that the root apex splits into two daughter roots. This is morphologically distinct from lateral root (LR) branching in the extant euphyllophytes, where LRs develop along the root axis at different distances from the apex. The process of root bifurcation is poorly understood, while such knowledge can be important, as it may represent an evolutionarily ancient strategy that roots recruited to form new stem cells or meristems. In this study, we examined root bifurcation in the lycophyte *Selaginella moellendorffii*. We characterized an *in vitro* developmental time-frame based on repetitive apex bifurcations, allowing us to sample different stages of dichotomous root branching and analyze the root meristem and root branching in *S. moellendorffii* at the microscopical and transcriptional level. Our results show that, in contrast to previous assumptions, initial cells in the root meristem are mostly not tetrahedral but rather show an irregular shape. Tracking down the early stages during root branching argues for the occurrence of a symmetric division of the single initial cell resulting in two apical stem cells allowing for root meristem bifurcation. Moreover, we generated a S. moellendorffii root branching transcriptome, which resulted in the delineation of a subset of core meristem genes. The occurrence of multiple meristem-related orthologues in this dataset, including inversely correlated expression profiles of a *SCARECROW (SCR)* versus a *RETINOBLASTOMA-RELATED1 (RBR1)* homologue suggests the presence of conserved pathways in the control of meristem and root stem cell establishment or maintenance.

**One-sentence summary:** The root of the spike moss *Selaginella moellendorffii* bifurcates following a symmetric cell division of the single stem cell and involves conserved genetic modules known from angiosperm roots.

---

Roots provide multiple key functions for plants, and their capacity to branch is important to anchor in a substrate and to forage for water and nutrients (Raven and Edwards, 2001; Motte et al., 2019). Root development and root branching have been in particular studied in angiosperms, e.g. Arabidopsis. Here, roots branch via the formation of lateral roots (LRs), which develop in the differentiated part of the root from pluripotent pericycle cells. After a series of well-orchestrated divisions, a new root meristem is formed that grows out of the parent root. Interestingly, important insights into the anatomical development and molecular regulation of LR formation were obtained by employing LR inducible systems, allowing synchronous induction of many LRs (Himanen et al., 2002; Himanen et al., 2004; Vanneste et al., 2005; De Smet et al., 2008; De Rybel et al., 2010; Jansen et al., 2013; Xuan et al., 2015; Crombez et al., 2016; Herrbach et al., 2017).

In contrast to angiosperms, Selaginella species, members of the lycophyte lineage, do not possess pluripotent cells that give rise to LRs (Fang et al., 2019). Instead, roots dichotomously branch at the tip through bifurcation of the apical meristem (Imaichi and Kato, 1989; Lu and Jernstedt, 1996; Otręba and Gola, 2011; Gola, 2014; Fang et al., 2019; Motte and Beeckman, 2019). Lycophytes were the first plants that developed roots (Hetherington and Dolan, 2018). Hence, studying their root system can contribute to the understanding of how root developmental mechanisms evolved. Lycophyte roots, however, originated independently from roots in the other land plant lineages (Hetherington and Dolan, 2019), but both molecular and anatomical data suggest a highly convergent evolution, and possibly even a common (partial) recruitment of a genetic program present in the rootless common ancestor of the different lineages (Huang and Schiefelbein, 2015; Motte and Beeckman, 2019). Based on fossil records, dichotomous root branching was found at the base of lycophyte evolution, putting it forward as an early innovation during the evolution of roots (Hao et al., 2010; Matsunaga and Tomescu, 2016; Hetherington and Dolan, 2017; Hetherington et al., 2020).

Unfortunately, despite intensive, but almost exclusively histological studies, the development and anatomy of the Selaginella root apical meristem and its bifurcation are still poorly understood. Multiple studies report the presence of one central tetrahedral initial cell (IC) or apical stem cell (Otręba and Gola, 2011 and refs therein) that, in analogy with the apical cell in the root meristem of certain ferns, was suggested to cut off daughter cells in four directions to provide cells for the root cap at the distal side, and cells for the growing root at the three proximal sides. In some ferns, it is clearly documented that the three proximal daughter cells each divide in an equal pattern, forming merophytes or clonally related groups of cells. Subsequent merophytes are superimposed to form different layers in the root (Hou and Hill, 2004). In contrast to ferns, such division pattern is less obvious in Selaginella roots. Furthermore, the branching process is problematic to study at the microscopical level. Live imaging is difficult due to the lack of biotechnological tools (Motte et al., 2020) and arbitrarily sampled roots rarely represent critical phases in which roots are undergoing bifurcation. Moreover, histological sectioning does not allow a high-throughput approach, and serial sectioning is required to obtain the relevant sections in which the IC is visible. As a consequence, the cell division events associated with root meristem bifurcation remain elusive, and different hypotheses still exist on how the doubling of the meristem might occur (Motte and Beeckman, 2019). One hypothesis suggests the disappearance of the IC due to segmentation followed by the recruitment of two new meristematic cells as two new ICs (Otręba and Gola, 2011). Barlow and Lück (2004), on the other hand, suggested that one of the daughter cells of the original IC becomes a new initial. Interestingly, in the Selaginella shoot meristem, a seemingly symmetric division of the apical cell initiates dichotomy (Jones and Drinnan, 2009). Such a symmetric longitudinal division of the apical cell is also the cause of thallus bifurcation in certain algae (Oltmanns, 1889; von Goebel, 1928; van den Hoek et al., 1995; Gola, 2014).

Being the first lycophyte with a sequenced genome, S. moellendorffii became an important model species for evolutionary research (Banks, 2009; Banks et al., 2011; Motte et al., 2020). Moreover, as root bifurcation seems to involve the initiation of a new root stem cell, dichotomous root branching could be used as a unique system to study root stem cell specification in an evolutionary context and to get insight into the complex process of dichotomy in general. Useful datasets and genetic tools start to become available, including several transcriptomic datasets from root samples (Ferrari et al., 2020; Motte et al., 2020; Fang et al., 2021). However, these studies are merely snapshots of the roots and do not allow to study temporal changes associated with the bifurcation process.

*S. moellendorffii* root branching cannot be induced by hormone treatments and a root branching inducible system, similar as for angiosperms, does not seem to be feasible (Fang et al., 2019), complicating the capturing of early bifurcating root meristems. Here, as an alternative, we developed a dichotomous branching time course assay to enrich for samples with bifurcating roots. We applied this assay for histological sectioning and whole-mount confocal microscopy. Based on this, and complementary to the current literature, we advocate that two new root meristems originate due to a symmetric division of the IC into two ICs. Furthermore, we exploited the assay to sample roots at regular intervals for RNA sequencing (RNA-seq) and revealed as such the *S. moellendorffii* root branching transcriptome. We show that transiently upregulated genes have a meristem signature, supporting the value of this dataset as a useful resource to identify candidate stem cell or meristem regulators in Selaginella. Moreover, a vast number of differentially expressed genes are homologous to Arabidopsis genes with a known role in stem cell or meristem functioning. This indicates, despite the structural difference in root meristems between Selaginella and Arabidopsis, a strong conservation in the pathways controlling meristem or stem cell functionality.

## RESULTS

### Developmental Time Frames of *S. moellendorffii* Root Bifurcation

The study of root bifurcation or branching in *S. moellendorffii* would take advantage of an inducible root branching system like the LR inducible system in Arabidopsis (Himanen et al., 2002). The added value of using such a system is that the initiation of new meristems can be synchronized and thus followed over time. A prerequisite is, however, to obtain a zero point at which no branching is ongoing. We previously reported that auxin transport inhibitors and a synthetic cytokinin were not useful to prevent root branching in *S. moellendorffii*, without affecting growth, and as such do not allow to obtain uniform starting material. Moreover, we did not succeed to identify a treatment that could induce meristem bifurcation itself (Fang et al., 2019). As a consequence, a synchronized root branching system that enables the study of root branching initiation does not seem to be feasible.

Therefore, we asked whether we could predict root branching based on the repetitive apex divisions taking place in *in vitro*-grown *S. moellendorffii* explants as an alternative approach. To characterize root branching, *S. moellendorffii* shoot explants were excised and transferred on fresh medium, which resulted in the spontaneous development of roots. First rhizophores were initiated, from which the root grew out (Fig. 1, A and B); subsequently, the roots repeatedly bifurcated dichotomously (Fig. 1, C and D; Supplemental Movie S1) (see also Fang et al., 2019). We followed the origin and bifurcation of in total 983 roots within three independent experiments, and observed all roots on a daily basis. Bifurcation events were macroscopically observed and recognized by the presence of two dome-shaped primordia at the root tip (Fig. 1C). In practice, it was difficult to exactly trace back the time of appearance for all rhizophores, as young rhizophores were almost colorless and mostly covered by the leaves. Still, taking the time point of the shoot explant transfer as the reference, we observed that a majority of the roots underwent the first bifurcation in a relatively short time span, more particularly, between 10 and 15 d post transfer of the explants (Fig. 1E).

**Figure 1.**
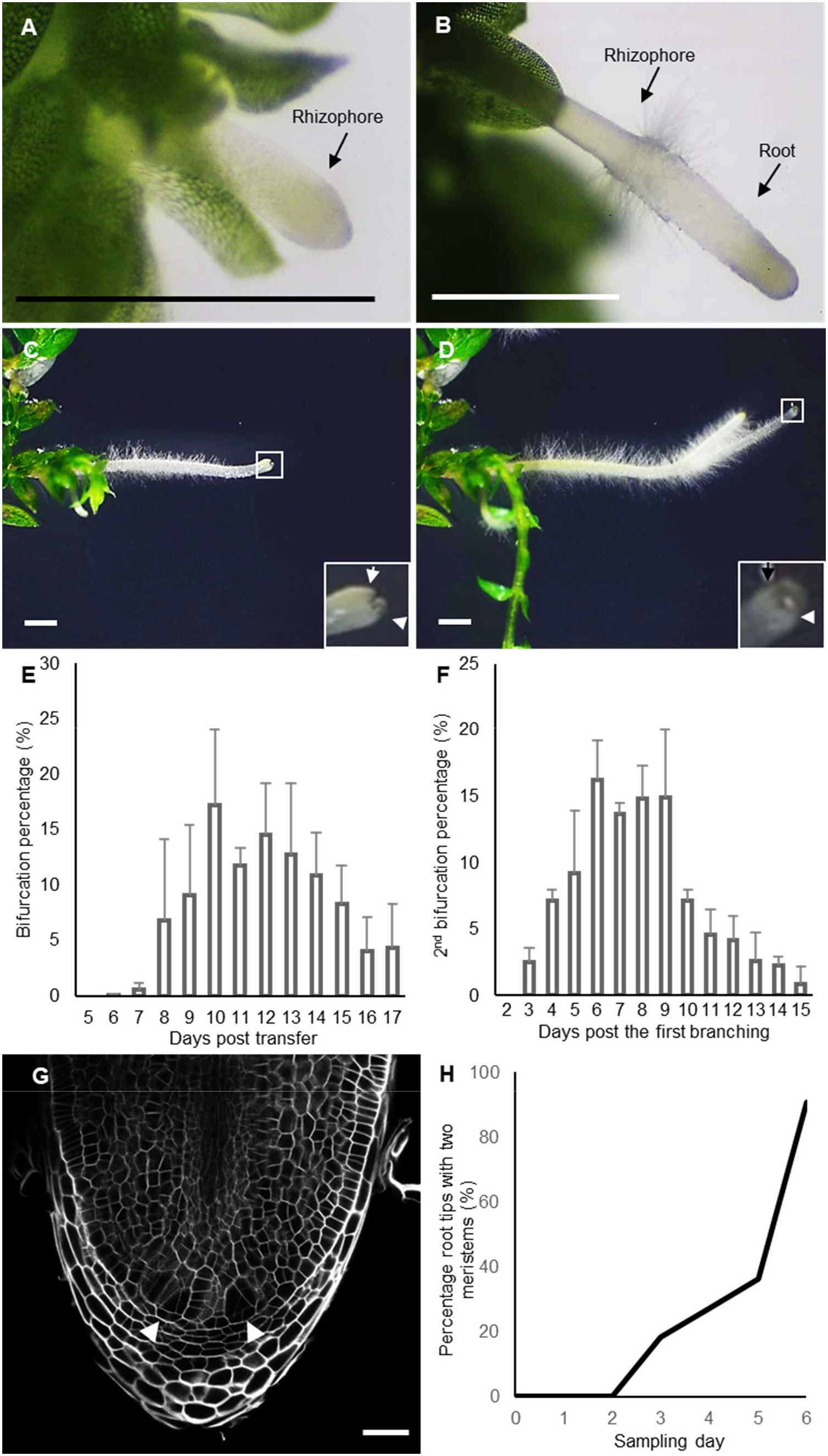
Dichotomous root branching in *Selaginella moellendorffii*. A, A young hairless rhizophore. B, Root emergence from the rhizophore. The transition from the rhizophore to the root is marked by the appearance of a collar of root hairs. Scale bars: 0.1 mm. C and D, Formation of first (C) and second (D) order bifurcated roots. The insets are magnifications of the squares. The arrows indicate newly formed root apices. Scale bars: 1 mm. E, Percentage of root tips that underwent the first bifurcation during 17 d after transfer on 1/2 MS. The bifurcation percentage was calculated as the number of bifurcated roots divided by the total number of roots. Values are averages of 3 repeats ± SD. F, Percentage of root tips that underwent the second bifurcation during 15 d after the first bifurcation.. The bifurcation percentage was calculated as the number of secondly bifurcated root apices divided by the total number of the apices from roots that underwent a first bifurcation. Values are averages of 2 repeats ± SD. G, Confocal image showing a bifurcated apical meristem: root tip with two new clearly recognizable young meristems (arrowheads). Scale bar: 50 μm. H, Percentage of bifurcated meristems in the root branching time course setup, i.e. days after the first branching. The percentage in Figure 1H was calculated as the number of root apices showing clear apical meristem bifurcation (see Fig. 1G) divided by the total number of sampled root tips. n (daily sample number) ≥ 10.

We continued to follow a subset of 361 root tips from two experimental repeats, and observed, taking the first bifurcation (Fig. 1C) as a reference point, a consistent short time frame for the second root bifurcation, which occurred in the majority of roots between 6 and 9 d after the first bifurcation (Fig. 1F). Hence, as the dichotomous branching seemed to occur according to a relatively regular periodicity, the branching time could be roughly predicted.

To study the timing of bifurcation in more detail at the histological level, we took the first bifurcation event as shown in Figure 1C as reference time point 0, and sampled at least ten root tips on a daily basis for microscopic analysis. All root tips were subjected to whole-mount confocal imaging to assess the appearance new meristems, as shown in Figure 1G. As previously shown, the apices at time point 0 never contained two apical meristems (Fang et al., 2019). Similarly, meristem bifurcation was not detected on the first and second day after the first branching (Fig. 1H). The first roots with two clear meristem regions were detected 3 d after the first branching. The number of roots with two meristem regions steadily increased over time and almost all root tips effectively bifurcated after 6 d (Fig. 1H).

Thus, in a relatively limited time frame after the first bifurcation event, practically all root tips bifurcated again. Because the presence of two meristems was visible from day 3 onwards, the onset of the bifurcation and related anatomical events could be anticipated to occur early during the time course. Furthermore, due to the steadily increasing percentage of bifurcated meristems, we could anticipate the enrichment of specific developmental stages during consecutive time points. Therefore, we employed this time course starting from newly branched roots to further characterize *S. moellendorffii* root branching, both from the histological and molecular point of view.

### Meristem Bifurcation Is the Result of a Symmetric Division of an Irregularly Shaped IC

To characterize the early events in *S. moellendorffii* root branching, we sampled root tips on different days after the first bifurcation event (time point 0) and subjected them to histological sectioning. At time point 0, two root tips are present each with an IC in the center of the meristem (Fig. 2, A and B). Within the meristems, merophytic cell clusters next to the IC with triangular shape could be easily recognized, illustrating how daughter cells have been cut off from different faces of the IC, and subsequently divided (Fig. 2B). Starting from two days after time point 0, we were able to recognize two similar-sized ICs next to or close to each other in different root tips (Fig. 2, C-G). This challenges the current hypotheses on the initiation of dichotomous root branching in Selaginella, and might point to a rather symmetric division of this IC as a first stage during root bifurcation. In particular Figure 2C shows a clear image of two separate basal derivatives from the presumptive ICs that form root cap cells. Figure 2D shows a similar image, but here, one of the two presumptive ICs has two nuclei and is finishing a cell division. Figure 2E and F show root apices with two presumptive ICs separated by only one cell, while Figure 2G shows a separation of the two ICs by more cells. Overall, this indicates that two new ICs originate after a symmetric division, and continue to develop two new root primordia.

**Figure 2.**
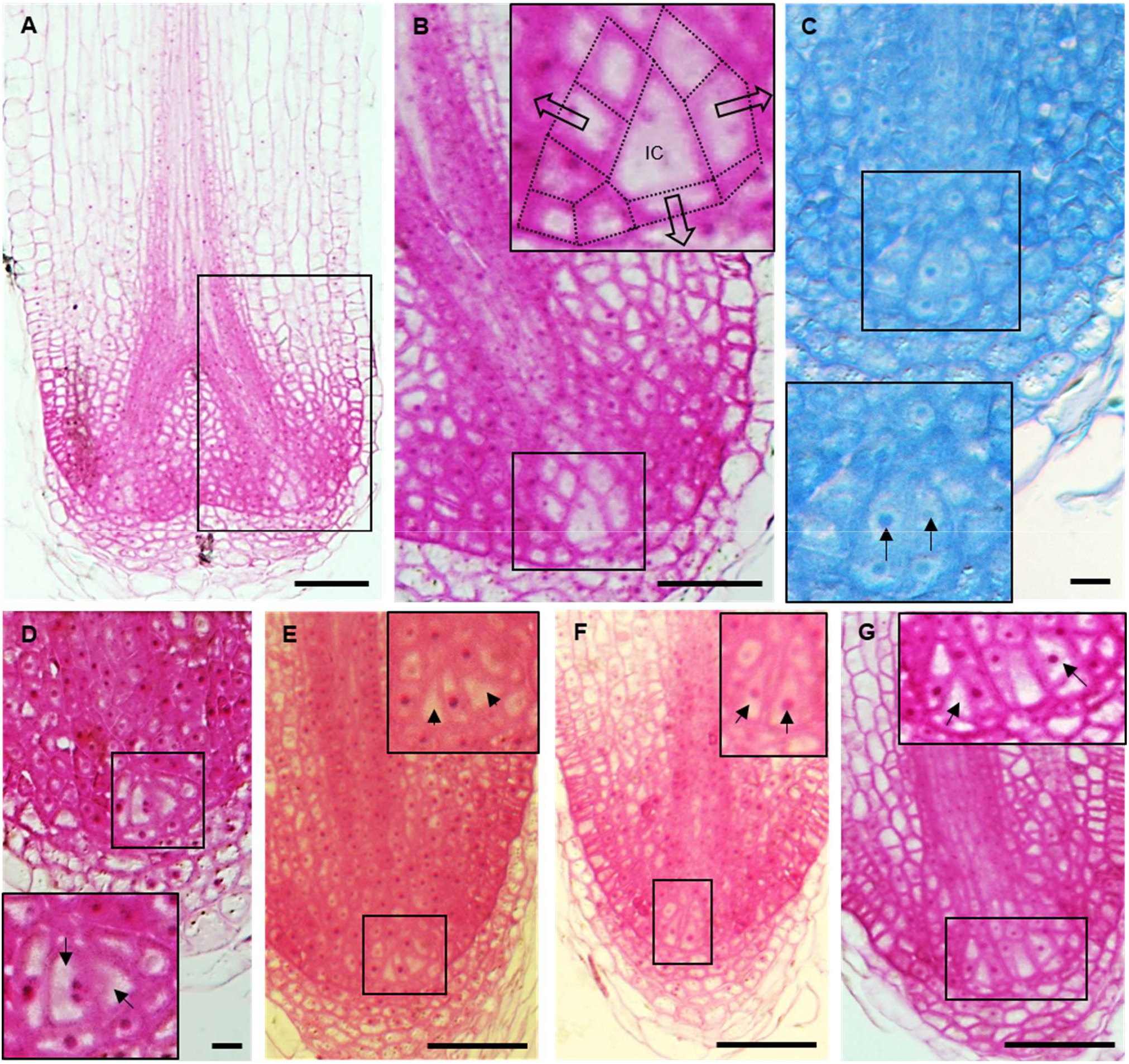
Histological sections from *S. moellendorffii* roots sampled during the root branching time course. A and B, A newly branched root tip containing two apical meristems sampled on day 0, (B) is a magnification of the indicated area in (A). The inset in (B) shows a magnification of the initial cell (IC) and its neighboring region. The initial and the presumptive most recent daughter cells are indicated by dotted lines. The hollow arrows indicate the expected directions in which the IC cuts off daughter cells from the different faces. C-G, Early-stage bifurcating meristems with two presumptive ICs next or close to each other. The insets shows a magnification of the indicated area, the arrows indicate the ICs. Scale bars: 50 μm, except for panel (C) and (D): 10 µm.

However, in many samples, we were not able to recognize such stages, and designation of the ICs was often not straightforward, possibly because ICs were regularly out of plane in the sections. Therefore, to confirm the observed and seemingly symmetric divisions of the IC, and to increase the workflow in order to analyze more samples at the same time, we decided to use the same approach by performing whole-mount confocal imaging. Supplemental Movie S2 shows a full stack of an unbranched root, and illustrates how this technique enables better designation of the IC by going through the sections. This stack was used to reconstruct the root (Fig. 3A) and clearly present the organization of the meristem. To have a better view on the structure of the IC, we first used whole stacks for 3D cell segmentation and isolated the IC. Earlier reports with two-dimensional histological sections hypothesized that the root IC is tetrahedral (Imaichi and Kato, 1989; Otręba and Gola, 2011). However, segmentation of the IC from several samples showed that its shape is not uniform and not tetrahedral (Fig. 3B; Supplemental Movie S3 to S7). Most side views, but not all, show triangular faces, while most distal faces are irregularly shaped, and are only triangular-like in a few cases. Hence, the IC in *S. moellendorffii* roots is in general not tetrahedral, and its irregular shape makes it difficult to be recognized, especially in sections that follows only one plane.

**Figure 3.**
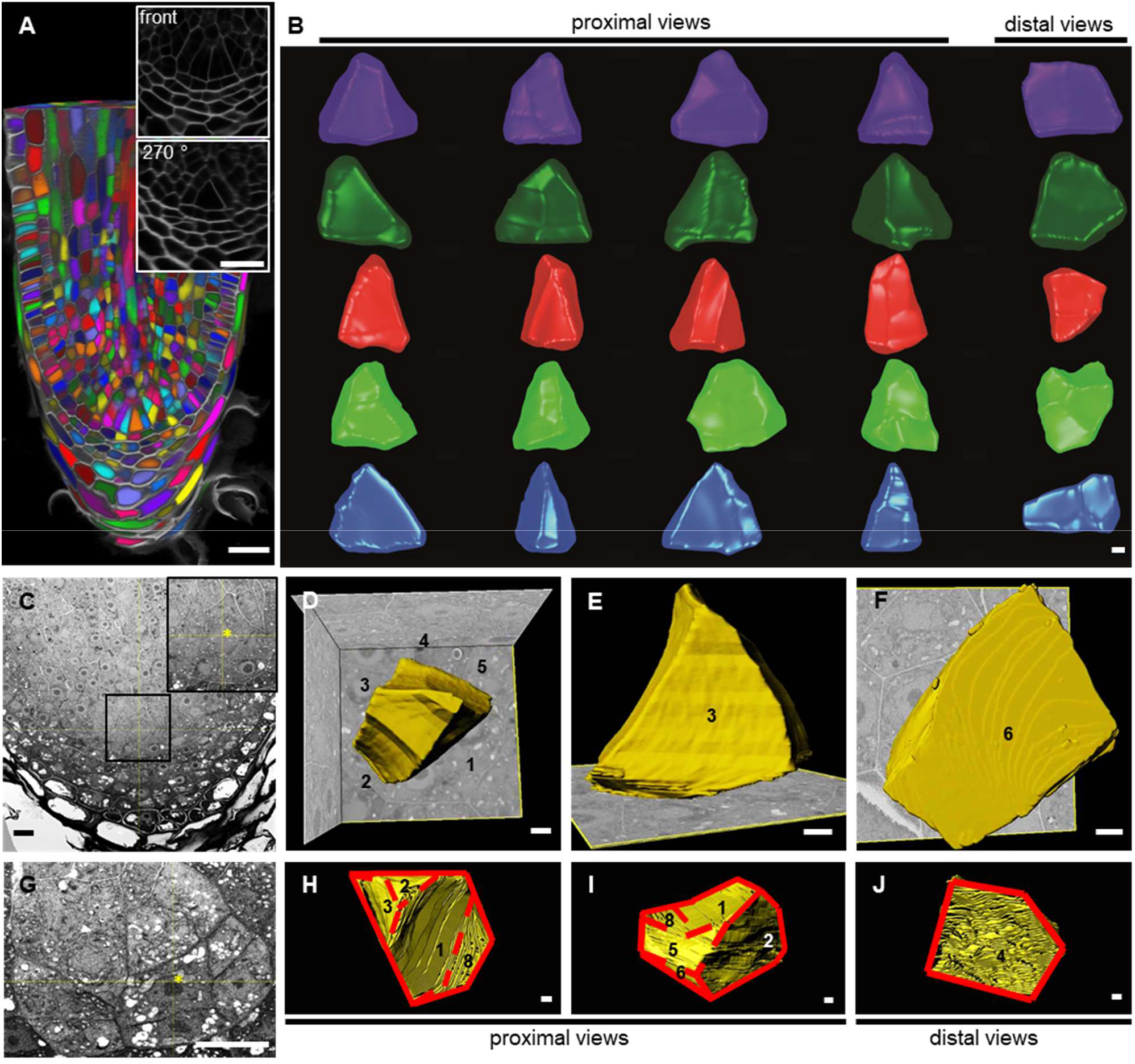
3D imaging to segment ICs. A, Example of a *S. moellendorffii* root generated from a whole-mount confocal stack (Supplemental Movie S2). Individual cells are randomly colored. Insets show magnifications of the front view or the 270 degree turned view. Scale bars: 20 µm. B, Different views of five ICs segmented from confocal stacks. The fifth IC was segmented from the root in panel A. See Supplemental Movies S3-S7 for a full overview of the ICs Scale bar: 2 µm. C and G, SBF-SEM longitudinal sections (xz-views) showing two ICs. The inset shows a magnification of the indicated area in (C). The asterisks indicate the ICs. Scale bars: 10 μm. D-F and H-J, Serial section reconstruction of the ICs. The numbers indicate the corresponding faces of the cells. See Supplemental Movies S8 and S9 for a full overview. The solid and dashed lines in (H-J) separate the different faces. Scale bars: 1 μm.

To confirm this irregular shape of the IC, we also subjected two root samples to serial block face scanning electron microscopy (SBF-SEM), allowing a much higher resolution of the cells and cell walls (Fig. 3, C-J; Supplemental Fig. S1; Supplemental Movie S8 and S9). The ICs seemed to have daughter cells at different faces, with cell plates or young cell walls seen in the mother and the most recent daughter cells (Fig. 3, C and G; Supplemental Fig. S1, A-B and F-G), which suggests that these cells were undergoing or had just finished cell divisions. 3D reconstruction based on the serial sections showed that one IC has five or six faces (face 4 and 5 might be considered as one, see Fig. 3D and Supplemental Fig. S1E), instead of four in a tetrahedron. Furthermore, it has a wedge-like shape with two opposite triangular faces (faces 1 and 3) and a rectangular bottom (face 6) (Fig. 3, D-E; Supplemental Fig. S1, C-E, Supplemental Movie S8). The second IC had three major proximal faces (faces 1-3), all with irregular triangular-like shapes, and a pentagonal distal face (face 4) (Fig. 3, H-J; Supplemental Fig. S1, H-J; Supplemental Movie S9). Hence, the ICs are irregular in shape, and not per se having a triangular face in all longitudinal planes. The IC might have this irregular shape as this cell is mitotically active (Fujinami et al., 2017), and therefore the shape changes during intensive cell divisions. This is an important observation to take into consideration when looking for ICs in sections, and explains why it is often troublesome to make a correct designation. By using whole stacks, it is possible to reorient the stack in such a way that ICs are clearly recognizable (as illustrated by the 270 degree view in Fig. 3A), as there always seems to be two faces with a triangular shape.

We next sampled root tips during the root branching time course in independent experiments covering different time points. The roots were subjected to whole-mount confocal imaging and stacks were made and segmented to reorient sections and to select planes with clear ICs (Fig. 4). Again, in some roots, two ICs of similar size were positioned next to each other (Fig. 4, A-C; Supplemental Fig. S2, A-C). This further supports the hypothesis for a symmetric division of the IC, which is most probably the first anatomical event leading to root meristem bifurcation.

**Figure 4.**
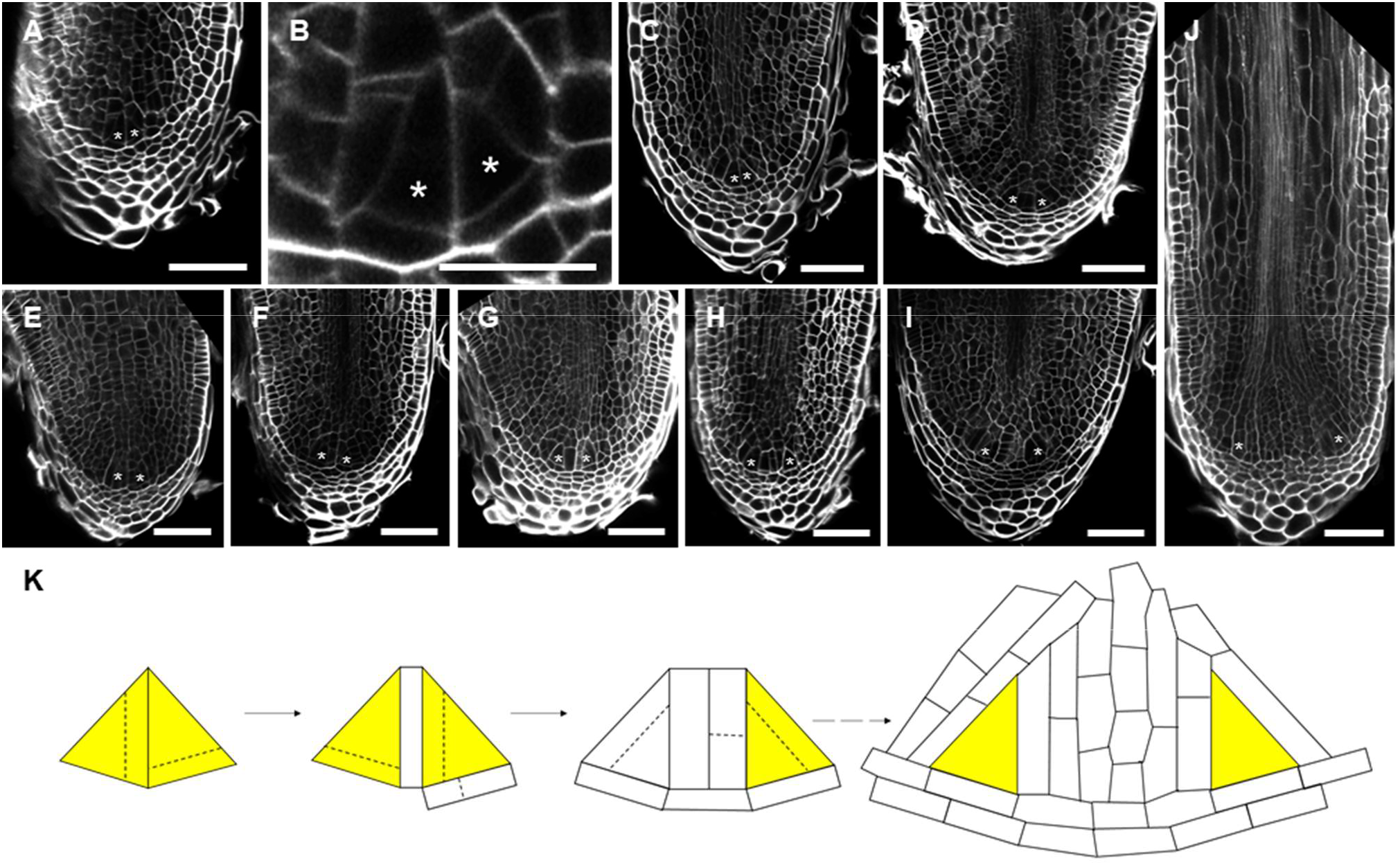
Early stages during *S. moellendorffii* root bifurcation. A-J, Confocal images of different stages during branching sampled between 1 and 5 days in the root branching time course. (B) shows a magnification of the root in (A). Scale bars are 50 µm, except for (B), where it is 20 µm. Asterisks indicate the location of the ICs. A-C, Two ICs are next to each other. See Supplemental Figure S2, A-C for more examples. E-G, Two ICs are only separated by one or two layers of cells. See Supplemental Figure S2, D-F for more examples. H-I, Further development of the two meristems, including enlargement of the root tip. K, Scheme showing the mechanism of root bifurcation as deduced from independent root samples during the early time points of root branching. Dashed lines indicate possible cell divisions.

In the absence of genetically encoded markers, live imaging to show that such division indeed results in the formation of two meristems is unfortunately not possible in *S. moellendorffii*. Still, in multiple samples at early time points, we could, based on the cellular pattern, clearly observe that each of the ICs produced their own distal daughter cells in the root cap (Fig. 4, A and B; Supplemental Fig. S2A), indicating that each of these two cells underwent at least one additional formative division to cut off a root cap cell. Furthermore, we also observed multiple root tips in which the two ICs were very close to each other and separated by only one or two layers of short cells (Fig. 4, D-G; Supplemental Fig. S2, D-F). These layers seemed to originate from daughter cells of the new ICs. Based on these observations, it is of high likelihood that the two ICs originally neighbored each other, supporting the hypothesis that dichotomous root branching in *S. moellendorffii* is a result of a symmetric division of the original IC. In this scenario, the two new ICs would each cut off daughter cells in different directions, forming two new meristems (Fig. 4K). These two new meristems or root primordia are initially still embedded in the macroscopically unbranched root, which radially expands because of the doubled meristem (Fig. 4, H-J), but later on grow out to form two branches as described before (Otręba and Gola, 2011).

### The *S. moellendorffii* Root Branching Transcriptome

#### Cyclic Gene Expression Profiles Support a Return to the Initial Meristem State

Based on the confocal imaging, almost 40% of the root tips formed two clearly distinguishable new meristems 5 d after the first branching (Fig. 1H), hence we supposed that programs to initiate dichotomous branching mainly occur before this time point. To identify possible molecular mechanisms with a role in root branching initiation in *S. moellendorffii*, we employed the developmental root branching assay for an RNA-seq experiment and sampled root tips on each day from 0 to 5 d after the first branching (time point 0), in which 300-µm apical parts were sampled to enrich for the meristematic region, while non-meristematic root regions were sampled separately (Fig. 5A).

**Figure 5.**
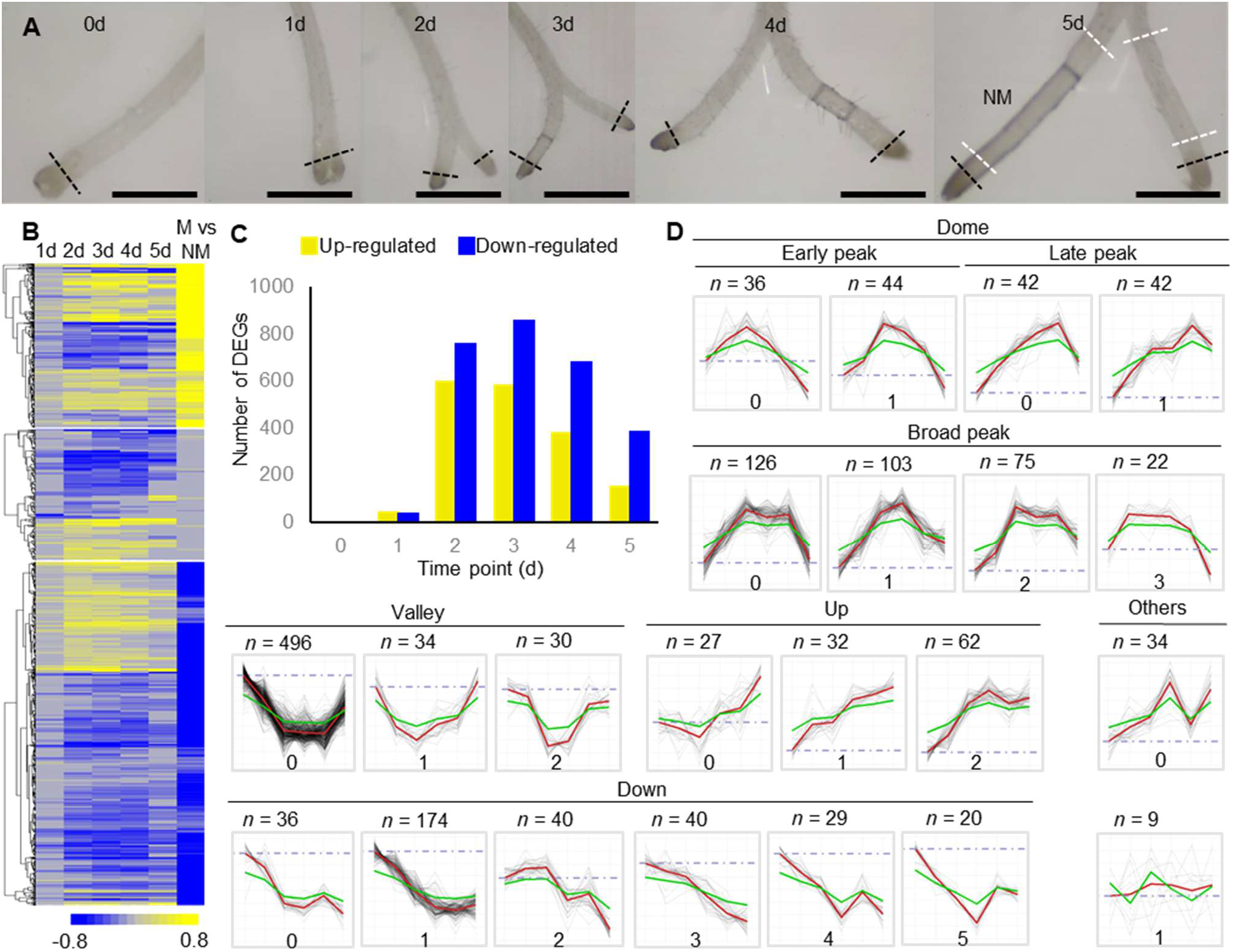
*S. moellendorffii* root branching transcriptome. A, Experimental set-up, showing sampled regions (dotted black lines: meristematic region; white: non-meristematic region (NM)). Scale bars are 1 mm. B, Heatmap showing expression [log_2_ fold change (FC)] of all root branching genes on the different days in the root branching course versus day 0, or for the meristematic (M) versus non-meristematic (NM) region. Gray values indicate no significance. C, Overview of numbers of up-regulated and downregulated differentially expressed genes (DEGs) at each time point during root branching. Note the decrease in number at the end of the time course, indicating a return to the initial state. D, Profiles of co-expression clusters based on weighted gene co-expression network analysis (WGCNA) of the root branching genes. Clusters were grouped according to their shapes: Dome and Valley groups – the transiently upregulated and downregulated clusters, respectively; Up and Down groups – constitutively upregulated and downregulated clusters, respectively. Based on the peak position and shape, the Dome group was further grouped into Early peak, Late peak and Broad peak subgroups. In each cluster, the red curve represents the average expression of all genes in the cluster, and the green curve represents the eigengene of the cluster.

Taking into account a fold change (FC) of at least 2 and an adjusted p-value (BH FDR corrected) of maximum 0.05, 6601 differentially expressed genes (DEGs) were retrieved between the meristematic and non-meristematic part. 3410 genes were higher expressed in the apical parts and were therefore designated as “meristem-enriched genes”. Within the branching time course, however, only 15 genes were differentially expressed with a |FC| > 2 (adjusted p-value ≤ 0.05; LRT). None of these DEGs had a straightforward annotation (Supplemental Table S1). This low number is not unexpected, as the different root branching samples only differed developmentally and, most likely, in a small region within the same tissue. Differences in gene expression may even only occur in one cell, and hence become diluted within the sampled tips. Still, 1553 genes were significantly (adjusted *p*-value ≤ 0.05; LRT) affected over the branching time course (Fig. 5, B and C; Supplemental Table S1). We therefore designated them provisionally as “root branching genes”. Many of these genes were differentially expressed at multiple time points (Fig. 5B; Supplemental Table S1), corroborating a relevant differential expression. Interestingly, the number of DEGs at first steadily increased during the time course and peaked on 2 and 3 d after the first branching, and then decreased towards the end of the time course (Fig. 5C). Hence, the final state showed less differential expression and was more similar to the original state.

To get a better insight into the expression profiles, the 1553 “root branching genes” were subjected to a cluster analysis via WGCNA (Langfelder and Horvath, 2008). The analysis revealed 22 main expression profiles (Fig. 5D), which we further grouped according to their shapes. As such, the clusters could be classified into four major groups: we designated the transiently upregulated and downregulated gene clusters as “Dome” (further grouped into Early peak, Late peak and Broad peak subgroups) and “Valley”, respectively; constitutively upregulated and downregulated clusters were named “Up” and “Down”, respectively; only a few genes could not be classified into any one of these groups (Fig. 5D). Overall, the majority of genes clustered in the Dome (490/1553) or the Valley (560/1553) groups, which fits the decreased number of DEGs towards the end of the time course.

In summary, both the number of DEGs per time point and the clustering of the expression profiles support a generally cyclic expression pattern, which might be correlated with developments at the cellular level: a starting state with a single meristem that rapidly initiates bifurcation, or evolves into an early bifurcation state and then evolves again into two separated single meristems resembling the initial state.

#### Root Branching Transcriptome to Identify Core Meristem-Enriched Genes

The Selaginella root branches in association with the development of two root meristems. Therefore, especially at an early stage with two meristems, we expected meristem genes to be differentially expressed: when the meristem bifurcates, transcripts of genes specifically expressed, or repressed, in the IC or in its neighborhood, would be relatively more, or less, abundantly present. Indeed, at an early stage, two young meristems are present within a non-expanded root tip. When the two meristems and the root tip enlarge (Fig. 4), this abundance would get diluted and evolve towards the end of the time course to a relative expression level as in a non-bifurcated meristem. Within the 1553 “root branching genes”, 452 out of the 3410 meristem-enriched genes were present (Supplemental Table S1), and hence, while this is still an overrepresentation (Fig. 6, A-C; Supplemental Table S2), a big portion of the meristem-enriched genes were not differentially expressed during meristem bifurcation. We presume that the group of “meristem-enriched genes” as such might still encompass many genes that do not associate with the meristem itself, but with another part of the root tip. The other possibility that these genes might be associated with maintaining meristem identity and therefore are not differentially expressed during bifurcation cannot be excluded. Still, especially meristem-enriched genes that are differentially expressed during the root branching time course (“root branching genes”) (Fig. 6, A and B), are expected to be associated with the root meristem function and might be of special interest.

**Figure 6.**
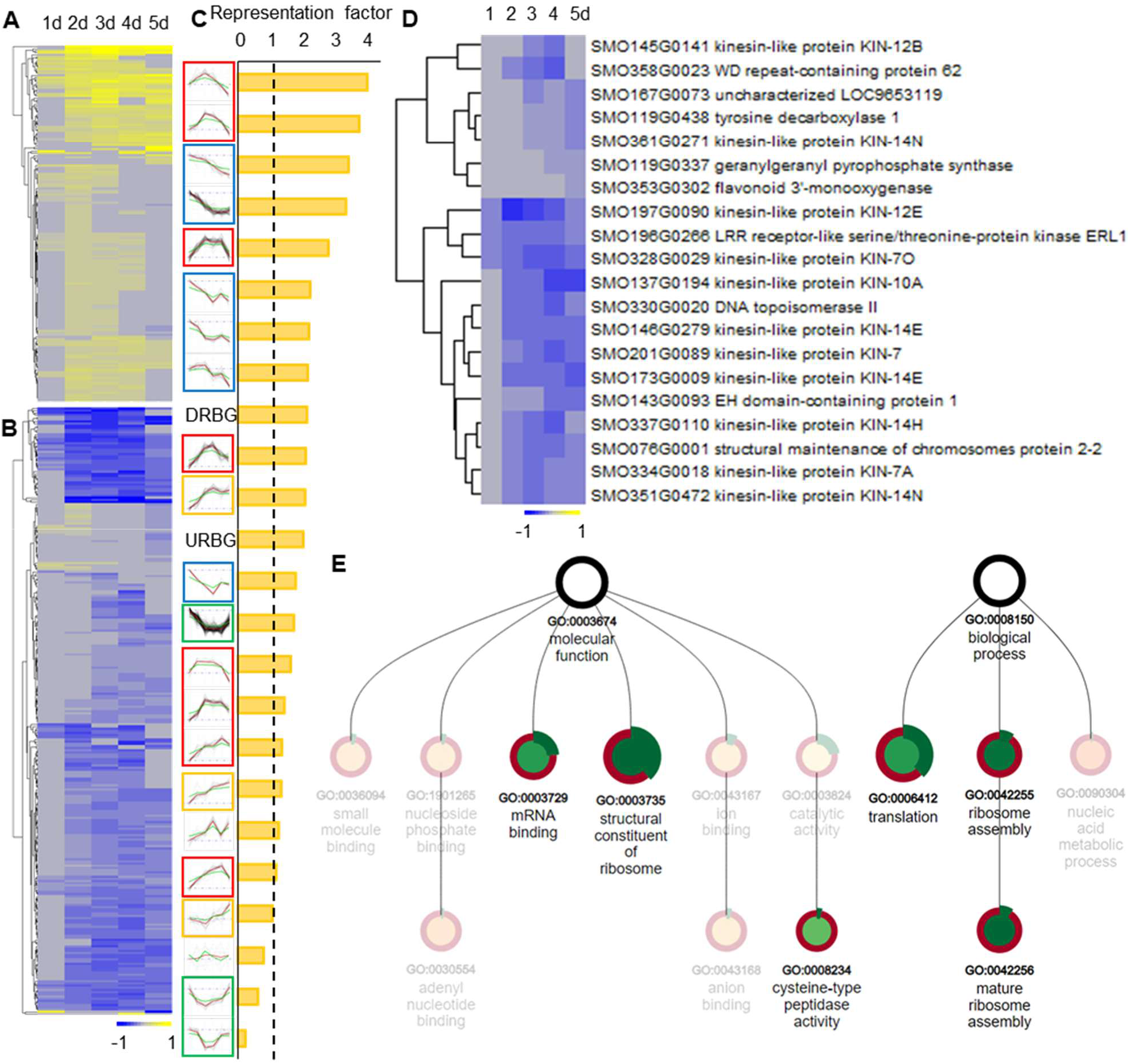
Overview of root branching meristem-enriched genes. A and B, The heatmaps show the expression [log_2_ fold change (FC)] of meristem-enriched genes upregulated (A) or downregulated (B) during root branching. C, Representation of the meristem-enriched genes in different expression clusters. The meristem-enriched genes were in particular overrepresented in transiently upregulated (Dome) clusters in red frames and constitutively downregulated (Down) clusters in blue frames. Orange and green frames indicate constitutively upregulated (Up) and transiently downregulated (Valley) clusters, respectively. DRBG, Downregulated root branching genes; URBG, Upregulated root branching genes. D, Heat map showing the expression [log_2_ fold change (FC)] the top 20 meristem-enriched genes that were downregulated during root branching. The descriptions of the genes were retrieved from Phytozome (JGI) or NCBI. E, Enrichment analysis of the meristem-enriched genes that were upregulated during root branching. The graph shows enriched Molecular Function and Biological Process Gene Ontology (GO) terms. Node size is scaled by the p-value of the enriched GO term, and node color (from green to red) is determined by the enrichment fold of the term. Negative enrichment folds are faded. The outer green band is determined by the subset ratio. The graph was generated by PLAZA version 4.5 (https://bioinformatics.psb.ugent.be/plaza/versions/plaza_v4_5_dicots/).

We were intrigued by the overrepresentation of meristem genes in the downregulated root branching DEGs, which showed a 1.9-fold overrepresentation in meristem genes (Fig. 6C), which could point to a loss of meristem identity or meristem activity during the bifurcation process. In the GO, InterPro and MapMan term enrichment analyses of this group, we noticed some terms related to microtubule-binding, cytoskeleton or cell division, including Heat Shock Protein 90-, Elongation Factor-, and kinesin-related terms (Krtková et al., 2012; Zhu and Dixit, 2012; Krtková et al., 2016), but also “microtubule binding”, “cytokinesis” and “cell cycle” (Supplemental Table S3). Indeed, a lot of cell cycle-related genes could be found in the top downregulated meristem-enriched genes, including extensin, callose synthase and phragmoplast-related *KINESIN-LIKE (KIN)* genes (Supplemental Table S1). When taking into account the top meristem-enriched genes in this subset, in particular KIN7, 10, 12 and 14 proteins were found (Fig. 6D), known to be associated with phragmoplast formation or orientation, and hence involved in cell plate formation and cell division (Zhu and Dixit, 2012). Thus, the group of downregulated meristem-genes were mainly related to cell division, which points to an arrest or inhibition of cell divisions during root branching. A more detailed analysis showed that these genes were especially present and enriched in Down clusters (137 genes) or in the big Valley-0 cluster (117 genes) (Fig. 6C; Supplemental Table S2). These clusters in general showed a middle to late downregulation, which is also notable on the general expression profile of these genes (Fig. 6, B and D). Possibly, cell divisions in the original meristem stopped or at least decreased, while daughter cells of the two new ICs took over the meristem activity towards the end of the time course.

The 173 of the meristem-enriched root branching genes that showed an upregulation in the root branching time course might be of interest. These genes were especially present and enriched in the early responsive (Early peak) Dome cluster, showing an expression profile that went up relatively early during the branching course, but returned to the basal level again at the end (Fig. 5D). Therefore, these clusters also show a much higher overrepresentation of meristem-enriched genes (Fig. 6C). The early upregulation of these genes indicates that two young meristems were formed early, which then matured towards the end of the branching time course.

To get insights into the function of the 173 meristem-enriched genes that were upregulated during the branching, GO, InterPro and MapMan term enrichment analyses were performed. These analyses clearly showed an enrichment in terms related to cell growth, and more particularly, in ribosome- and translation-related terms (Fig. 6E; Supplemental Table S3). The gene set indeed included a broad range of ribosomal proteins (Supplemental Table S1). Interestingly, such genes or terms are typically enriched in young meristematic angiosperm root cells as well. For example, Arabidopsis single-cell transcriptomic studies showed that ribosomal protein-encoding genes are enriched in meristematic clusters (Ryu et al., 2019; Shulse et al., 2019; Zhang et al., 2019). Hence, the 173 genes seemed to be associated with young meristematic cells, underlining and validating the value of the root branching dataset to select core-meristem genes.

Thus, this set can be used to highlight genes with a putative role in the *S. moellendorffii* root meristem, especially as the subset of 173 genes are probably differentially expressed in the young meristem, possibly even in the IC, but not in the root cap or other regions of the root tip.

We queried the 173 genes for orthologs of genes encoding better characterized proteins in Arabidopsis. This subset contains, besides many ribosomal proteins, an *ARABIDOPSIS RESPONSE REGULATOR15 (ARR15)* ortholog (SMO274G0055) and an *AUXIN RESPONSE FACTOR7 (ARF7)* ortholog (SMO361G0080). *ARR15*, negatively regulating cytokinin signaling, is important for stem cell specification in the Arabidopsis root meristem (Müller and Sheen, 2008), and the auxin-induced *ARF7* is important during LR initiation, leading to a new root meristem (Okushima et al., 2005). Their presence in this set of genes suggests a possible importance of the two genes in *S. moellendorffii* root meristem establishment or maintenance as well, and as such a, possibly partially, in the conserved genetic program of root stem cell specification.

#### Candidate Regulators of Dichotomous Root Branching

To further identify possible regulators in the root branching process, we queried all 1553 root branching genes for transcription factors and orthologs of genes with well-described roles in *Arabidopsis* root meristem maintenance or LR branching, including auxin and cytokinin signaling genes, receptor kinases and cell cycle genes (Supplemental Table S5). Transcription factors were collected via PlantTFDB v5 (Tian et al., 2019). As we noticed that not all ARF and ARR transcription factors were recognized in the PlantTFDB, we also looked for their orthologs and included them in the transcription factor query (Supplemental Table S5).

Among these possible regulators, 40 were differentially expressed during the root branching time course (Fig. 7A), and might be involved in the regulation of dichotomous root branching. These included four auxin-responsive *ARF* genes, two cytokinin-regulating *ARR* genes, and orthologs of the auxin transporter *PIN2*, the cytokinin-responsive *CYTOKININ RESPONSE FACTOR1 (CRF1)*, the cytokinin receptor *AHK2*, the gibberellic acid signaling gene *REPRESSOR OF GA-LIKE1 (RGL1)*, the ethylene signaling gene *ETHYLENE-INSENSITIVE3-LIKE1 (EIL1)* and the brassinosteroid signaling gene *BES1/BZR1 HOMOLOGUE4 (BEH4)*. The presence of these genes might indicate a role of hormonal signaling in a very complex way in dichotomous root branching and the associated meristem maintenance.

**Figure 7.**
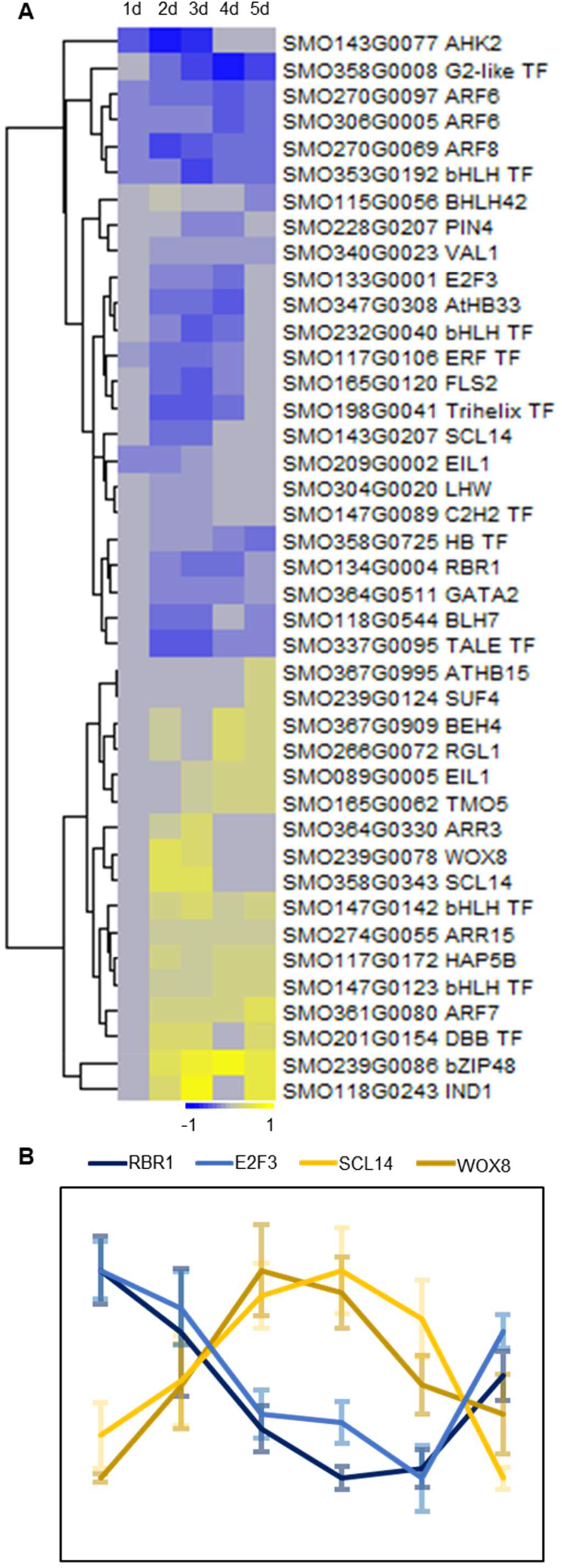
Expression profiles of the transcription factors encoding DEGs and other genes with their Arabidopsis homologs known to be involved in root meristem control or lateral root (LR) formation. A, Heatmap showing the expression [log_2_ fold change (FC)] profiles. The description shows the best Arabidopsis hit according to Banks et al. (2011) or, if not present, the transcription factor type was retrieved according to Plant Transcription Factor Database (PlantTFDB) v5. B, Relative expression of homologs of *RETINOBLASTOMA-RELATED1 (RBR1), E2F TRANSCRIPTION FACTOR-3 (E2F3), SCARECROW (SCR) (SCR-LIKE14 (SCL14))*, and the *WUSCHEL RELATED HOMEOBOX (WOX)* family *(WOX8)*. Error bars indicate SE (n = 4).

We also noticed the presence of homologs of both *TARGET OF MONPTEROS5 (TMO5)* and *LONESOME HIGHWAY (LHW)*. In Arabidopsis, the two form a dimer and are together crucial for vasculature initiation (De Rybel et al., 2013; De Rybel et al., 2014). This control is shown to be highly conserved in all vascular plants (Lu et al., 2020). In Arabidopsis, the two transcription factors have a different spatial expression pattern and overlap at the vascular domain (De Rybel et al., 2013). Within the *S. moellendorffii* root branching course, *LHW* was early downregulated, but reached its basal level again at the end, whereas *TMO5* was upregulated in particular at the end (Fig. 7A). Hence, the *LHW* and *TMO5* homologs showed a common high expression level towards to the end of our time course (from 3 and 5 d), when two young meristems were present, possibly indicating increased vascular development at that stage.

Interestingly, only a few transcription factor-encoding DEGs showed an early upregulation. In particular, a *WUSCHEL RELATED HOMEOBOX (WOX)* (SMO239G0078) and a *SCARECROW (SCR)* (SMO358G0343) homolog, both important and transcriptionally interacting stem cell regulators (Shimotohno et al., 2018; Motte et al., 2020), showed a strong early induction (Fig. 7A) and are highly co-expressed (Fig. 7B). Possibly, these were involved in the control of dichotomous root branching. Interestingly, also *RBR1* (SMO134G0004), important in stem cell fate decision as well and a negative interactor of SCR (Cruz-Ramírez et al., 2012), was found in the list. As shown for Arabidopsis, *RBR1* interacts with E2F transcription factors (Wildwater et al., 2005), of which a homolog, SMO133G0001 was also differentially expressed in our dataset. Noteworthy, at least in Arabidopsis, *RBR1* and *E2F3* show a high degree of co-expression (Magyar et al., 2012), which we also observed in the *S. moellendorffii* root branching time course (Fig. 7B).

In contrast to the Dome expression profile of the *WOX* and *SCR* homologs, we observed a downregulating Valley profile for the *S. moellendorffii* RBR1 and E2F3 homologs. The two group of genes thus showed an inversely correlated expression pattern (Fig. 7B), which intriguingly fits within the pathway model proposed for Arabidopsis.

## DISCUSSION

Lycophyte roots branch dichotomously due to bifurcation of the apical meristem. Activities of the IC have been assumed to play a key role in this process, especially in branching initiation, whereas it remained unclear how its cellular activities are associated with root bifurcation in the lycophyte *S. moellendorffii*. Previous reports on Selaginella root bifurcation suggested that the single root apical IC undergoes segmentation and becomes indistinguishable, which is followed by the initiation of two new ICs at the flanks and the formation of two new root meristems (Imaichi and Kato, 1989; Otręba and Gola, 2011). Barlow and Lück (2004) hypothesized that, after a set of rigid cell divisions, a derivative of the IC becomes a new IC to form the second root meristem. In contrast, our results point out that root bifurcation starts with a symmetric division of the original IC, which gives rise to two new ICs. Such a symmetric division to initiate dichotomous root branching in Selaginella has never been reported. Additionally, it was stated that a longitudinal symmetric division of a tetrahedral cell is unlikely to occur, because the surface area of the inserted cell wall is relatively big and would be inefficient (Cooke and Paolillo Jr, 1980; Gola, 2014). However, we showed that the IC is irregular and possibly changing in shape, and in general not tetrahedral. As such, ICs might acquire geometries that are better suited to undergo a symmetric division.

Still, there could be other factors controlling the IC to undergo the symmetric division to initiate dichotomous root branching. As the stem cell is surrounded by multiple cells in the meristem, interactions and feedbacks from the neighboring cells may also contribute to the first division initiating root branching in land plants. In Arabidopsis, the auxin reflux from surrounding endodermal cells allows auxin accumulation in the founder cells for transition to LR initiation (Dubrovsky et al., 2008; Marhavý et al., 2013; Cavallari et al., 2021), also showing a critical role of auxin in the control of the first asymmetric division. However, we detected differential downregulation of multiple auxin-responsive *ARF* genes in the branching time course, suggesting a reduction in the auxin response during the process. Thus, auxin is not likely to accumulate in the IC to induce the first symmetric division. Supportive for this, we earlier found that Selaginella root branching is not induced by auxins (Fang et al., 2019). Moreover, a spatial accommodation by neighboring cells, which may involve mechanical forces, is also essential for Arabidopsis LR initiation. Specifically, pericycle cells swell before the first asymmetric division, which is actively accommodated by the neighbor endodermis (Vermeer et al., 2014; Vermeer and Geldner, 2015; Ramakrishna et al., 2019). However, it is totally unknown whether the IC and the neighboring derivatives need to undergo any morpho-cytological alteration to promote root branching initiation in Selaginella.

During Arabidopsis root growth, (lateral) root branching formation takes place periodically: protoxylem cells in the basal meristem undergo temporal oscillations in auxin concentrations and auxin-responsive gene expression levels, which allows these cells to transport these priming signals to the pericycle cells, which also get primed, forming competent sites to initiate LRs (De Smet et al., 2007; Moreno-Risueno et al., 2010; Van Norman et al., 2014; Xuan et al., 2016; Laskowski and ten Tusscher, 2017). In Selaginella, roots also branch repetitively but dichotomously (Fang et al., 2019), which coincides with a generally cyclic gene expression profile. Possibly, this involves priming signals as well, which may be included in our dataset.

It has been hypothesized that lycophyte roots have a shoot origin (Gensel et al., 2001), which should result in shared developmental mechanisms between the shoot and root meristem. This was for instance shown in the lycophyte Lycopodium, where bifurcation of both meristems starts with the emergence of mitotically active cells in a similar quiescent region (Fujinami et al., 2021). In contrast, Selaginella bifurcation of the shoot meristem begins with a symmetric division of the IC (Christopher and Andrew, 2009).This fits our observation of the same division in the root, and corroborates the shoot origin of the root in lycophytes. It also underlines the possible differences in root anatomy and bifurcation mechanisms, as not all lycophyte subclades have single ICs, and supports along with the fossil record that roots originated multiple times during the evolution of lycophytes (Gensel et al., 2001; Fujinami et al., 2017; Fujinami et al., 2020; Hetherington et al., 2021).

Although roots originated multiple times during evolution, it is hypothesized based on comparative transcriptomics studies that a common rootless ancestor of vascular plants possessed an inheritable genetic tool kit that descendants recruited for similar root meristem-related biological processes, resulting in a highly conserved root developmental program within vascular plants (Huang and Schiefelbein, 2015; Ferrari et al., 2020). Details on this conserved genetic program are lacking, but using a root branching time course in *S. moellendorffii*, we succeeded to introduce specific core-meristem genes and showed the presence of multiple homologs of Arabidopsis genes with a known role in meristem establishment. This further corroborates that the genetic control of root development, despite morphological differences, is indeed highly conserved and further supports the parallel recruitment of an, at least partial, common genetic program.

As the organization of the Selaginella root meristem is less complex than that of seed plants, a more elementary pathway controlling stem cell fate decisions may be presumed (Motte et al., 2020). Thus, it might be anticipated that Selaginella requires a more limited set of genetic players to install and maintain the stem cell. We identified homologs of some known root stem cell controllers in Arabidopsis, including, *WOX, SCR, RBR1 and E2F3*-family members, as the candidate regulators of dichotomous root branching in Selaginella. Moreover, we found that their expression profiles fit within the pathway model in Arabidopsis. Hence, conserved pathways, involving SCR-dependent regulation of a WOX gene and a RBR1-SCR feedback loop for stem cell fate decisions, or at least roles of RBR1 and E2F3 similar as those of the Arabidopsis root (Wildwater et al., 2005; Stahl et al., 2009; Cruz-Ramírez et al., 2012), could be speculated. It is important to note that WOX functioning in angiosperm stem cell control and its action via transcriptional repression is restricted to the WUS subclade (Dolzblasz et al., 2016; Zhou et al., 2018), which is not present in *S. moellendorffii* (Wu et al., 2019). Thus, the presence of *WOX* as a branching DEG may imply a different mechanism, possibly as a transcriptional activator (Motte et al., 2020). It is tempting to hypothesize that these genes are the central components in the evolutionarily early pathways controlling the IC, and were recruited for both dichotomous and LR branching formation.

Overall, we generated a relevant dataset, in which multiple candidate regulators of root meristem bifurcation in *S. moellendorffii* could be identified that possibly show high conservation with similar pathways in seed plants. In contrast to existing *S. moellendorffii* datasets, which mainly focused on tissue-specific expression, we here characterized expression patterns during a developmental process in which new (root) stem cells are established. Although the molecular research using *S. moellendorffii* is still at its infancy, this dataset can be a useful resource to identify possible root meristem- and stem cell-related Selaginella or lycophyte genes. Due to the pioneering evolutionary position of this lineage during plant colonization of land, this resource can be very useful to identify conserved and early (root) stem cell regulators, and to unravel the early pathways that were at the base of pluripotency.

## MATERIALS AND METHODS

### Plant Materials

*S. moellendorffii* plants were obtained from the lab of Jo Ann Banks, Purdue University. Plants were grown aseptically under a 16-h light/8-h dark photoperiod at 20.25–43.2 μmol/m^2^/s in a 24°C growth chamber and roots developed after the transfer of explants to fresh medium as previously reported (Fang et al., 2019).

### Microscopy

To track root branching events, roots were observed daily starting from their appearance on rhizophores using a Leica S9I stereomicroscope.

For whole-mount confocal imaging, *S. moellendorffii* root tips were first fixed in 50% methanol and 10% acetic acid, and then subjected to modified pseudo-Schiff propidium iodide (mPS-PI) staining described by Truernit et al. (2008). Confocal imaging was performed using a Zeiss LSM 5 Exciter microscope (25x water immersion objective lens) with an argon ion laser at 488/505 nm as the excitation/emission wavelengths, or a Leica SP2 confocal microscope [63x water corrected objective (NA 1.2) lens and pinhole = 1 AU] with an argon ion laser at 514/600-650 nm as the excitation/emission wavelengths. Alternatively, modified staining was performed with a small change in the composition of chloral hydrate solution [8 g chloral hydrate, 3 mL glycerol, and 1 mL water (8:3:1)]. Z-stacks were taken for all samples to allow collective detection of the meristematic region present at different planes. A subset of samples, including the ones used for segmentation, were after fixation subjected to amylase treatment in order to digest starch granules. Root tips were washed three times with water and subsequently treated overnight with α-amylase at a concentration of 300 U/ml in a buffer containing 20 mM phosphate pH 7.0, 2mM NaCl, 0.25 mM Ca_2_Cl. Samples were then washed with water and fixed again in 50% methanol and 10% acetic acid for 1 h. Samples were subsequently washed two times with 1X PBS and transferred to ClearSee solution (Kurihara et al., 2015) and gently stirred for 1 week. Finally, the samples were stained for 1 h in ClearSee solution containing 0.1% Calcofluor White and were washed afterwards for 30 min in ClearSee solution before imaging. Confocal imaging of the Calcofluor White stained roots was conducted using 405 nm excitation and 425-475 nm wavelength detection using a Leica SP8X system. Confocal stacks generated for segmentation were imaged using a 63X water-corrected objective (NA of 1.2) and a pinhole of 0.6 AU. Stacks were further processed using MorphoGraphX (Barbier de Reuille et al., 2015) to identify ICs and to obtain clear sections showing the meristematic organization.

For histological sections, root tip samples were fixed in 1% glutaraldehyde and 4% paraformaldehyde, followed by dehydration and embedding in Technovit 7100 resin (Heraeus Kulzer) according to the manufacturer’s instruction. Transparent strips were used to orient tissues for proper orientation (Beeckman and Viane, 2000). 2-µm sample sections were cut using a Leica Reichert-Jung 2040 Autocut Microtome, dried on object glasses using a hot plate (20–40 °C), stained for cell walls with 0.05% toluidine blue or ruthenium red (Acros Organics) for 10 min and then rinsed with water to wash off excess dyes. The sections were then mounted in Depex medium (Sigma) and covered with cover slips for observation and photography using an Olympus BX53 microscope. For serial block face scanning electron microscopy (SBF-SEM), samples were fixed using 2% paraformaldehyde and 2.5% glutaraldehyde in 0.1 M Sodium Cacodylate pH 7.4. Next, samples were processed en-bloc using a ROTOTO staining as described previously (Fendrych et al., 2014). Finally, resin-embedded root tips were mounted on aluminum specimen pins (Melotte) using conductive epoxy (Circuit Works), and coated with 5 nm of Pt in a Quorum sputter coater. Next, samples were placed in the Gatan 3View 2XP in a Zeiss Merlin SEM, for imaging at 1.6 kV using the Gatan Digiscan II BSED detector. The system was set up to automatically take 1000 images at a pixel-size of 15 nm, cutting 100-nm sections of the block-face between each image, resulting in a volume of 150 x 150 x 100 µm.

For registration of the 3D image stack, IMOD (http://bio3d.colorado.edu/imod/) was used. Orthogonal views were obtained in Fiji (http://fiji.sc/Fiji). Segmentation was done using Microscopy Image browser (http://mib.helsinki.fi/) and rendering was done using Imaris (Bitplane).

### Extraction of Initial Cell Geometry

Extraction of initial cell geometry was performed using MorphoGraphX (Barbier de Reuille et al., 2015). Confocal images were prepared for segmentation using a Gaussian blur of 0.5 in the X, Y and Z directions. Segmentation was performed using the ITK watershed auto seeded algorithm with a threshold of 800 and segmentation errors were manually corrected. 3D cell meshes were constructed for the region of the root containing the IC, using the process marching cubes 3D with a cube size of 0.5, and 50 smooth passes at the time of mesh generation to minimize alterations to the mesh structure (Bassel, 2015). All other cells in the mesh were then deleted to isolate the IC, which was subject to a further 5 smooth passes using the smooth mesh tool to facilitate qualitative visualization.

### RNA-Seq

Sampled roots in the time course (300 root tips per sample, 4 biological repeats per time point) were cut off using a pair of stainless microscissors and then stored in RNAlater^®^ RNA Stabilization Solution (Thermo Fisher Scientific) according to the manufacturer’s instruction. Subsequently, microdissection was performed to collect root apices of 0.3 mm long, as well as non-meristematic regions between apices and branching point for the 5 d post first branching samples. The samples were then collected in 2-mL RNase-free Eppendorf tubes with an RNase-free metal ball [diameter: 5 mm, sprayed by RNaseZap™ RNase Decontamination Solution (Invitrogen™, Thermo Fisher Scientific)] and frozen in liquid nitrogen. After collection, samples were stored at −70 °C.

For RNA extraction, samples were ground three times using a QIAGEN Retsch Tissuelyser (30 Hz, 30 s). RNA extraction was done via TRIzol™ reagent (Thermo Fisher Scientific) and the RNeasy Plant Mini Kit (Qiagen) according to the provided protocols.

RNA concentration and purity were determined spectrophotometrically using a Nanodrop ND-1000 (Nanodrop Technologies) and RNA integrity was assessed using a Bioanalyzer 2100 (Agilent). Per sample, an amount of 250 ng of total RNA was used as input. Using the Illumina TruSeq Stranded mRNA Sample Prep Kit (protocol version: Part # 15031047 Rev. E - October 2013), poly-A containing mRNA molecules were purified from the total RNA input using poly-T oligo-attached magnetic beads. In a reverse transcription reaction using random primers, RNA was converted into first strand cDNA and subsequently converted into double-stranded cDNA in a second strand cDNA synthesis reaction using DNA PolymeraseI and RNAse H. The cDNA fragments were extended with a single ‘A’ base to the 3’ ends of the blunt-ended cDNA fragments after which multiple indexing adapters were ligated introducing different barcodes for each sample. Finally, enrichment PCR was carried out to enrich those DNA fragments that have adapter molecules on both ends and to amplify the amount of DNA in the library. Sequence-libraries of each sample were equimolarly pooled and sequenced on Illumina HiSeq 4000 (SBS 300 cycles, Paired End Reads:151-8-8-151) at the VIB Nucleomics Core (www.nucleomics.be).

### Expression Analysis

We used two versions of *S. moellendorffii* annotated genomes: the NCBI version (Annotation Release 100, released in 2018, containing 33,283 genes) and the JGI version (v1.0, released in 2007, containing 22,285 genes). While the former is the most recent, and based on transcriptome data more correctly mapped, the JGI version is more widely used in bioinformatic tools. Therefore, we performed the mapping/alignment based (i) on NCBI if there was a 1:1 BLAST hit between NCBI and JGI annotations (and including the corresponding JGI identifiers), and (ii) on JGI for genes that were uniquely present in the JGI annotation. Counts were quantified using Salmon (Patro et al., 2017) and normalization was done using the DESeq2 R package (Love et al., 2014). To correct for the batch effect, Limma (Ritchie et al., 2015) was applied.

Differential analyses of the count data were performed using DESeq2 (Love et al., 2014), only considering normalized read counts ≥ 10. PCA was used to identify and omit outliers within the biological repeats and differential gene expression was analysed using a multiple factor analysis (MFA).The expression during branching was assessed by a likelihood-ratio test (LRT) (Supplemental Table S1). Differentially expressed genes were selected based on an adjusted p-value [Benjamini & Hochberg (BH) (Benjamini and Hochberg, 1995) false discovery rate (FDR) corrected] ≤ 0.05. In addition, meristem DEGs were only retained in case of a |FC| ≥ 2. Heatmaps showing expression profiles were generated using the R package “pheatmap”. Weighted gene co-expression network analysis (WGCNA) was performed according to Langfelder and Horvath (2008) in the pipeline of DESeq2.

Statistical significance of the overlap between two groups of genes was calculated via the Fisher exact test. Representation factors were calculated by dividing the number of overlapping genes by the number of expected genes in the overlap.

Data for Gene Ontology (GO), InterPro and Mapman term enrichment were computed using the ‘Workbench’ function of the Dicots PLAZA tool version 4.5: https://bioinformatics.psb.ugent.be/plaza/versions/plaza_v4_5_dicots/ (Vandepoele et al., 2013; Van Bel et al., 2018), with default settings.

Homology or orthology between Arabidopsis and *S. moellendorffii* genes was based on the gene families in the Dicots PLAZA version 4.5.

RNA-seq data are available in the ArrayExpress database (https://www.ebi.ac.uk/arrayexpress) under the accession number XXXX.

## Supplemental Data

The following supplemental materials are available.

Supplemental Figure S1. Additional SBF-SEM images and proximal views of the reconstructed root ICs in *S. moellendorffii*.

Supplemental Figure S2. Additional confocal images taken during the root branching assay in *S. moellendorffii*.

Supplemental Movie S1. *S. moellendorffii* shoot explant with root formation that repetitive bifurcation of the roots.

Supplemental Movie S2. Confocal stack of an unbrached *S. moellendorffii* root.

Supplemental Movie S3. First IC in Figure 3B.

Supplemental Movie S4. Second IC in Figure 3B.

Supplemental Movie S5. Third IC in Figure 3B.

Supplemental Movie S6. Fourth IC in Figure 3B.

Supplemental Movie S7. Fifth IC in Figure 3B.

Supplemental Movie S8. IC in Figure 3D-F.

Supplemental Movie S9. IC in Figure 3H-J.

Supplemental Table S1: Processed RNA-Seq gene expression values.

Supplemental Table S2: Overlap of genes between different subsets or clusters and the meristem-enriched genes.

Supplemental Table S3: GO, Mapman and Interpro terms associated with the meristem-enriched genes that were downregulated in the root branching course.

Supplemental Table S4: GO, Mapman and Interpro terms associated with the meristem-enriched genes that were upregulated in the root branching course.

Supplemental Table S5: Selaginella orthologs or homologs from lateral root or root stem cell related gene families as retrieved from PLAZA version 4.5 (https://bioinformatics.psb.ugent.be/plaza/versions/plaza_v4_5_dicots/).

## ACKNOWLEDGMENTS

We thank Lieven Sterck and Ignacio Eguinoa for support in bioinformatic analyses, Anna Kremer, Benjamin Pavie, Peter Borghgraef and Saskia Lippens for assistance and guidance with the SBF-SEM, Davy Opdenacker for technical support and Annick Bleys for help in preparing the mansucript. We also would like to thank Moritz Nowack, Edyta Gola and Koen Geuten for the helpful discussions and comments on the manuscript.

## Supplemental data

**Supplemental Figure S1.**
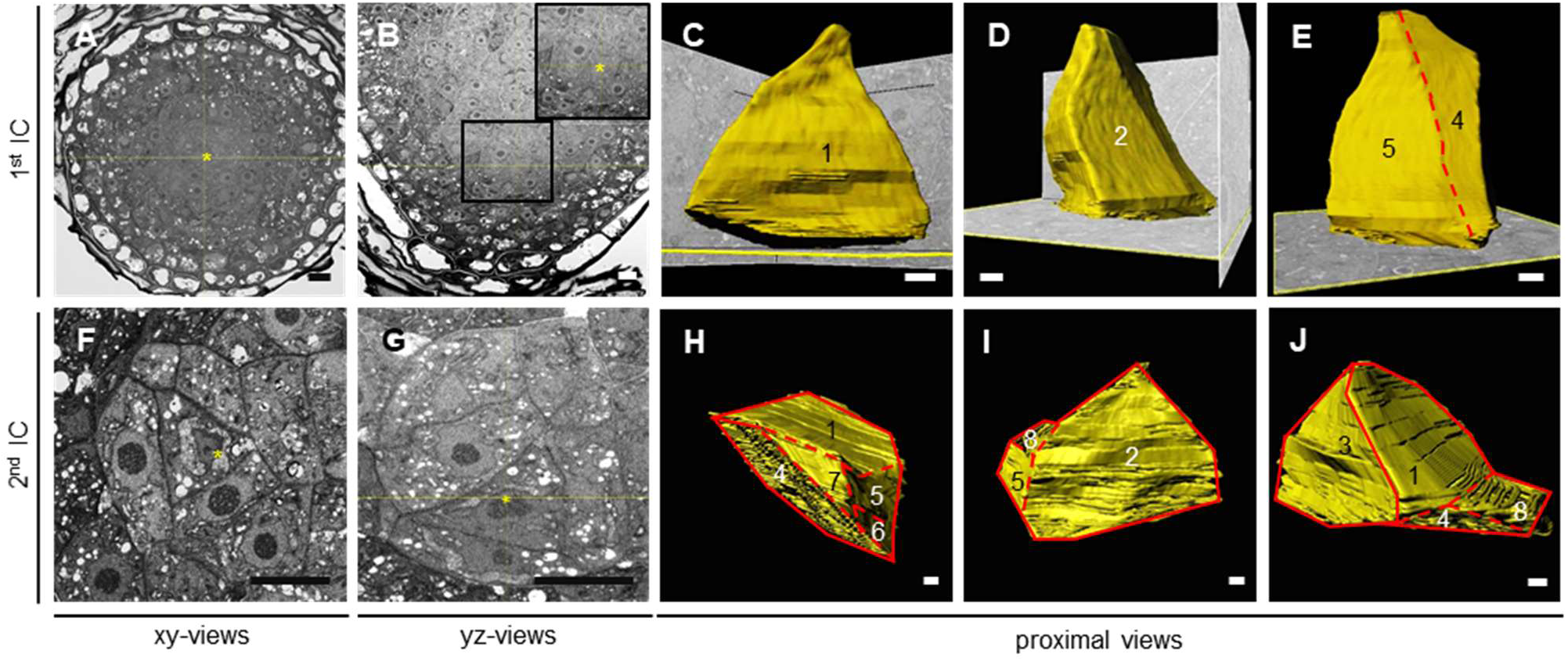
Additional SBF-SEM images and proximal views of the reconstructed root ICs in *S. moellendorffii*. (A-E) show the first IC and (F-J) show the second. (A and F) and (B and G) are transverse (xy) and longitudinal (yz) SBF-SEM sections, respectively. The inset shows a magnification of the indicated area in (B). The asterisks indicate the ICs. Scale bars: 10 μm. (C-E) and (H-J) show other proximal views of the reconstructed ICs, demonstrating that the ICs are not tetrahedral. The numbers indicate the corresponding faces of the cells. The solid and dashed lines in (H-J) indicate the main lines and the ones relatively difficult to be recognized, repectively. Scale bars: 1 μm.

**Supplemental Figure S2.**
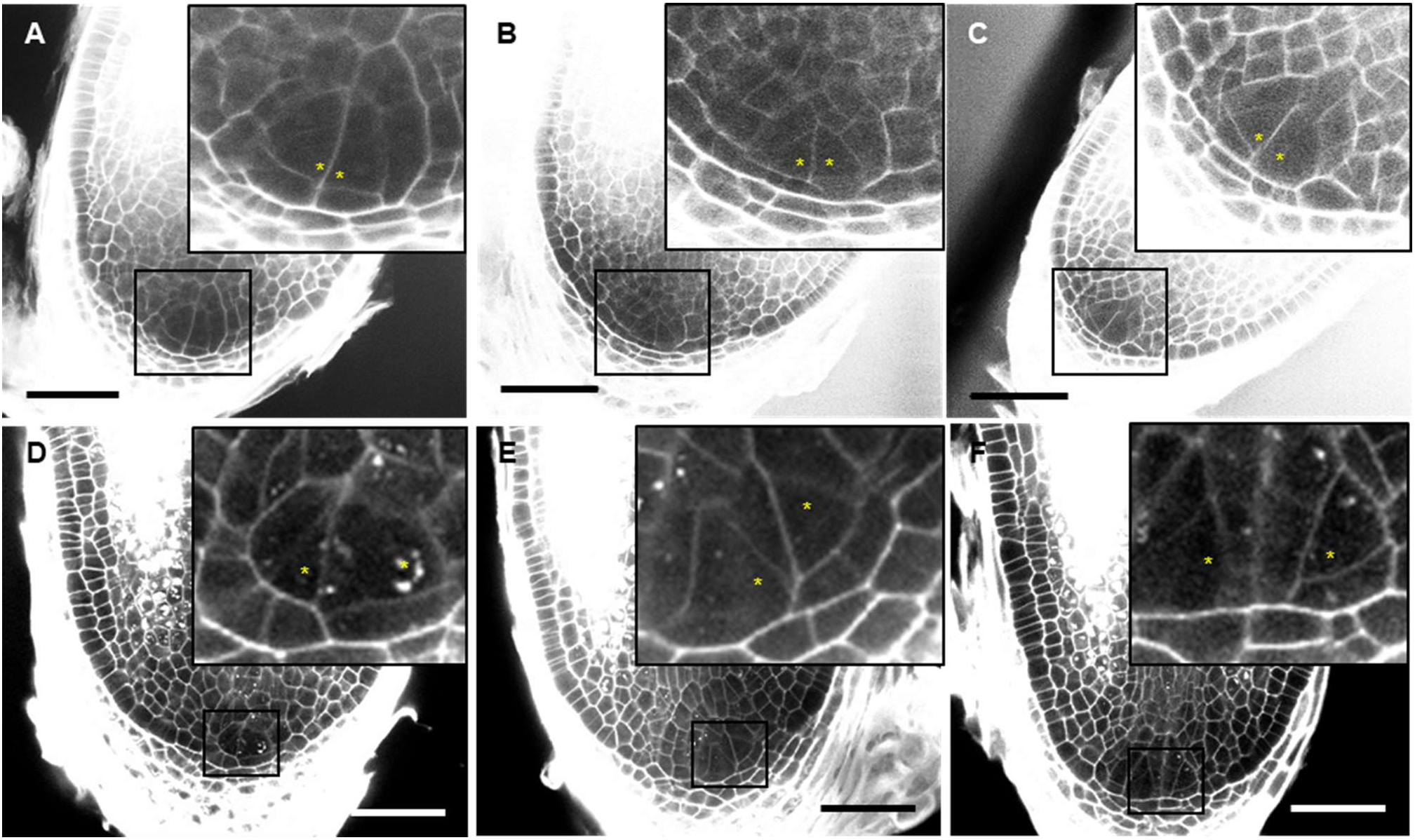
Additional confocal images taken during the root branching assay in *S. moellendorffii*. A-C, More examples of symmetric division of the original IC resulting in two new initials. D-F, More examples showing two initials that are only separated by one or two layers of cells, further supporting that they originated next to each other. Scale bars: 50 μm.

